# Per-residue optimisation of protein structures: Rapid alternative to optimisation with constrained alpha carbons

**DOI:** 10.1101/2025.11.24.690085

**Authors:** Ondřej Schindler, Tomáš Svoboda, Gabriela Bučeková, Radka Svobodová

**Affiliations:** National Centre for Biomolecular Research, Masaryk University, Kamenice 753/5, Brno, 625 00, Czech Republic; CEITEC – Central European Institute of Technology, Masaryk University, Kamenice 753/5, Brno, 625 00, Czech Republic

**Author notes:** Contributing authors.

**Keywords:** protein, residue, structure optimisation, force fields, GFN-FF, AlphaFold DB

## Abstract

In recent years, the number of known protein structures has increased significantly. Predictive algorithms and experimental methods provide the positions of protein residues relative to each other with high accuracy. However, the local quality of the protein structure, including bond lengths, angles, and positions of individual atoms, often lacks the same level of precision. For this reason, protein structures are usually optimised by a force field prior to their application in further research sensitive to structural quality. Protein structure optimisation, however, is computationally challenging.

In this paper, we introduce a general method *Per-residue optimisation of protein structures: Rapid alternative to optimisation with constrained alpha carbons* (PROPTIMUS RAPHAN). Rather than optimising the entire protein structure at once, PROPTIMUS RAPHAN divides the structure into overlapping residual substructures and optimises each substructure individually. This approach results in computational time that scales linearly with the size of the structure. Additionally, we present PROPTIMUS RAPHAN_GFN-FF_, a reference implementation of our method employing a generic, almost QM-accurate force field, GFN-FF.

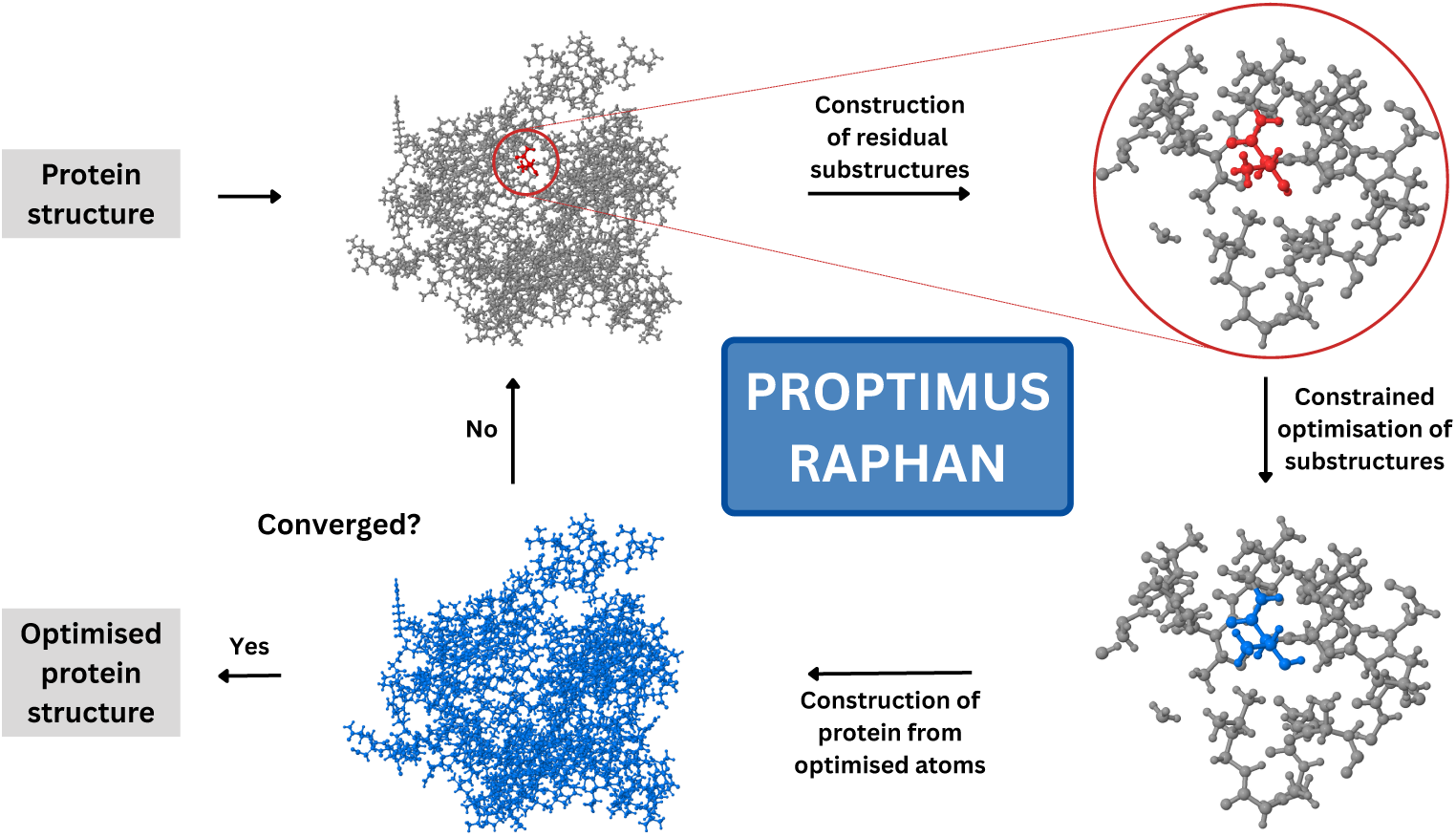

We tested PROPTIMUS RAPHAN_GFN-FF_ on 461 AlphaFold DB structures and demonstrated that our approach achieves results comparable to the optimisation of the structure with constrained alpha carbons in significantly less time.

**Scientific Contribution:** The main contribution of this work is the PROPTIMUS RAPHAN method and its reference parallelisable implementation PROPTIMUS RAPHAN_GFN-FF_. Because the time requirement increases linearly with the size of the structure, PROPTIMUS RAPHAN_GFN-FF_ optimises on average 5 000 atoms per hour and a common CPU. Therefore, prior to any research sensitive to protein structure quality, our method can be employed to obtain protein structures closer to QM-accuracy.

## 1 Introduction

Proteins are a fundamental part of all living organisms and are extensively studied for their crucial roles in many biological functions [1]. The 3D structure of a protein, the arrangement of atoms in space, is the crucial factor that determines the function of a protein [2]. In recent years, the number of predicted and experimentally determined protein structures has increased significantly. The Protein Data Bank [3] currently contains more than 240 thousand experimentally determined protein structures. On the other hand, more than 200 million predicted structures are available in the AlphaFold DB [4, 5], and over 700 million structures are stored in the ESM Metagenomic Atlas [6]. Furthermore, several freely available web services are available to predict the structure of a protein sequence, with the latest ones being based on artificial intelligence [7–9].

Machine learning methods aim to accurately determine the positions of alpha carbons in a protein and thus the relative positions of individual amino acid residues. Current machine learning approaches can predict the positions of alpha carbons in a protein with accuracy comparable to experimental methods [10]. In contrast, detailed characteristics of amino acid side chains, such as bond lengths, bond angles and residue interactions, are not predicted with the same precision [11, 12]. A similar phenomenon is also observed in experimentally determined protein structures, where the local quality of the structure could be limited by the accuracy of the experiment (e.g., resolution in the case of X-ray crystallography).

The local quality of protein structure is essential for correct results in various applications of computational chemistry [13]. Applications sensitive to structure quality are, for example, docking [12], QSPR modelling [14], partial atomic charge calculations [14], or calculations using semiempirical or QM/MM methods [15]. Therefore, predicted protein structures are usually optimised before conducting the main research [13, 16].

Force field methods are commonly used for protein optimisation. These methods replace the electronic structure of the molecule used in quantum chemistry methods with classical mechanical interatomic interaction potentials, resulting in a fast calculation [17, 18]. However, even with the use of a force field, optimisation of proteins with more than ten thousand atoms remains computationally challenging [17]. Therefore, various constraints can be introduced during protein structure optimisation to reduce the number of degrees of freedom and accelerate computation. Constraining alpha carbons at a specific location in the space during optimisation is one of the most frequently used techniques [4, 13, 16]. This approach is particularly suitable for structures predicted by machine learning methods, as alpha carbons are the most accurately predicted atoms. However, even with these constraints, force field methods still retain their quadratic computational complexity in relation to the number of atoms for single-point energy calculations and, consequently, for the overall protein structure optimisation [17].

In this work, we introduce a general iterative method Per-residue optimisation of protein structures: Rapid alternative to optimisation with constrained alpha carbons (PROPTIMUS RAPHAN). In every iteration of the PROPTIMUS RAPHAN method, a substructure is created for each protein residue, which includes the residue and its neighbouring residues. These substructures are then optimised separately. Then, the optimised substructures are used to reconstruct a partially optimised protein structure. This approach allows for linear computational dependence on protein size. The method is applicable to any force field. In our work, we also establish a reference implementation of PROPTIMUS RAPHAN method employing a force field, GFN-FF (PROPTIMUS RAPHAN_GFN-FF_) [17].

## 2 Methods and implementation

The PROPTIMUS RAPHAN method is developed based on the Cover approach, a divide-and-conquer algorithm, which has already been successfully utilised multiple times to calculate partial atomic charges [19–21]. The Cover approach is an approximation that neglects long-range weak interactions, thereby allowing the protein to be partitioned into smaller fragments. The PROPTIMUS RAPHAN divides the protein into overlapping residual substructures for each residue, and substructures are then optimised separately. The time required to optimise one substructure is approximately constant, as the substructures are similar in size. This leads to approximate linear computational complexity with respect to the number of atoms. Moreover, the use of the divide-and-conquer algorithm ensures high parallelisability.

We developed a reference implementation of PROPTIMUS RAPHAN in Python. Before the PROPTIMUS RAPHAN computation begins, the protein structure is loaded using the BioPython library [22]. We refer to this BioPython object in the following text as the *target structure*, because it tracks and stores all intermediate results during the optimisation. Moreover, a set of *optimised atoms* is defined for each protein residue. *Optimised atoms* are all atoms of the residue except for the alpha carbon and the N-H peptide bond atoms. Instead, the N-H peptide bond atoms from the following residue are included in the *optimised atoms*.

PROPTIMUS RAPHAN is an iterative method. Each iteration consists of the following steps (see Figure 1):

1. **Construction of residual substructures:** A residual substructure is created for each non-converged *target structure* residue. The residual substructure includes all atoms that are within 6 Å to at least one *optimised atom* of this residue. Moreover, a minimum set of other atoms is included to ensure that only bonds between two carbon atoms are cut [21]. Atoms of each residual substructure that do not belong to the *optimised atoms* of the specific residue are divided into two groups:

- *Flexible atoms* are all atoms that are closer than 4 AÅ to at least one *optimised atom* and are not alpha carbons.
- *Constrained atoms* are the remaining atoms of the residual substructure that belong to neither the *optimised* nor the *flexible* atoms. We implemented this step using the BioPython [22] and RDKIT [23] libraries.

2. **Constrained optimisation of residual substructures:** All residual substructures undergo their own optimisation. During optimisation, the coordinates of *constrained atoms* cannot be altered and therefore only positions of *flexible atoms* and *optimised atoms* are optimised. Residual substructures are optimised by GFN-FF [17] provided within the xtb software [24]. For that reason, we refer to our implementation as PROPTIMUS RAPHAN_GFN-FF_. Solvation is included in the calculation by using the implicit solvent model ALPB [25]. The number of GFN-FF optimisation steps is limited to prevent extensive changes in atom coordinates. Specifically, the maximum number of GFN-FF optimisation steps is determined by the sum of *optimised atoms* of the residual substructure and the current PROPTIMUS RAPHAN_GFN-FF_ iteration index.
3. **Construction of protein from optimised atoms:** The *target structure* coordinates are updated based on the corresponding optimised atomic positions.

**Fig. 1.**
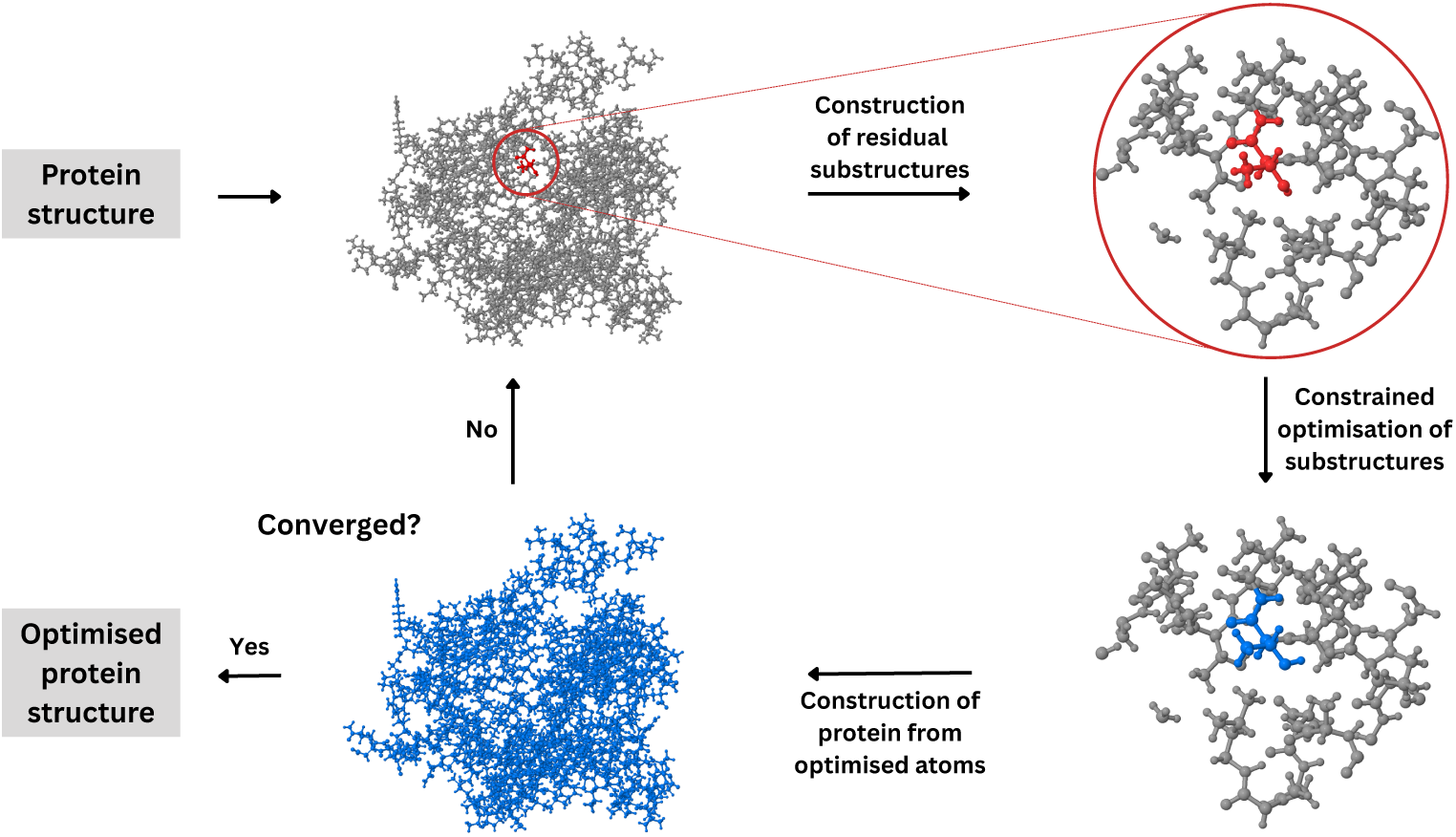
Scheme of the PROPTIMUS RAPHAN method.

An advantage of PROPTIMUS RAPHAN is the ability to continuously exclude already converged residues (i.e., residues whose position no longer changes during optimisation) from the iterative process. This feature significantly speeds up the entire calculation.

To increase accuracy, the entire iterative process is repeated twice. In the second run, 8 AÅ is used instead of 6 AÅ when creating substructures. In addition, all atoms of the residue except for the alpha carbon are considered to be *optimised atoms* of the residue.

For the purposes of testing and reproducibility, we also included in the PROPTIMUS RAPHAN_GFN-FF_implementation the option to directly run the GFN-FF optimisation with constrained alpha carbons (GFN-FF_C_*_α_*). In addition, we provide Python scripts for comparing protein structures using the Biopython [22], RDKit [23], Biotite [26], and MDAnalysis [27] libraries, which were used in Section 3. The PROPTIMUS RAPHAN_GFN-FF_ implementation, alongside the scripts used for comparison, is provided in the supplementary material.

## 3 Results and Discussion

To compare the accuracy and speed of PROPTIMUS RAPHAN_GFN-FF_ with those of the original method GFN-FF_C_*_α_*, we designed a series of experiments and selected a test dataset.

### 3.1 Datasets

The testing dataset would contain selected protein structures from AlphaFold DB [5]. In particular, the sample came from the fourth version (v4) of predictions for Swiss-Prot sequences [28]. The selection was performed in such a way that exactly one random structure was selected from each interval of 10 on the size scale ranging from 200 to 5 000 heavy atoms. Consequently, the dataset contained 480 structures. Then it was necessary to furnish the structures with hydrogens, since no predictions from AlphaFold DB included them. For this, we employed the tool PB2PQR [29] with the PROPKA [30, 31] titration state method. For each structure, the pH was randomly selected from a range of 0 to 14. This was the original dataset, and hereafter we will refer to it as SET_ORIG_. Having the SET_ORIG_ prepared, three different optimisation procedures were applied to it (see Figure 2). The following procedures took place:

- Optimisation using GFN-FF_C_*_α_* method. The subjected set of structures will be further denoted as SET_C_*_α_*.
- Optimisation using PROPTIMUS RAPHAN_GFN-FF_ method. The subjected set of structures will be further denoted as SET_RAPHAN_.
- Optimisation using PROPTIMUS RAPHAN_GFN-FF_ method and a subsequent one using the GFN-FF_C_*_α_* method. The subjected set of structures will be further denoted as SET_RAPHAN+C_*_α_*.

**Fig. 2.**
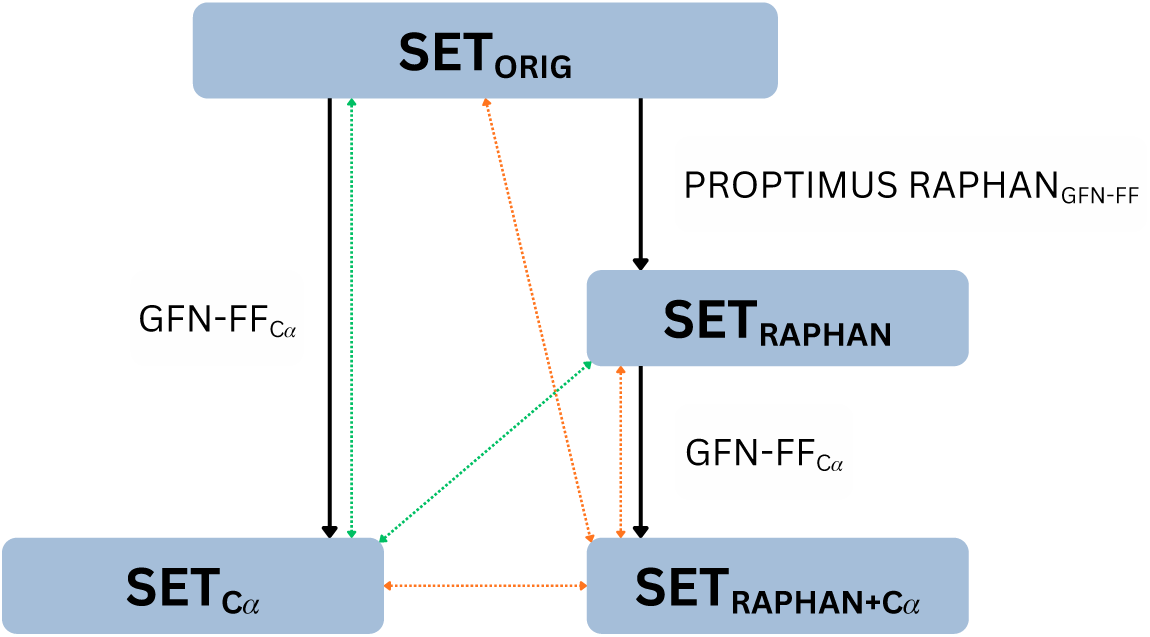
Schematic diagram describing the relationships between datasets. Black lines represent optimisations. Dotted lines represent the pairs of datasets compared as described in Sections 4 (green lines) and 4.1 (orange lines).

All optimisations were launched within our implementation. All optimisations were performed on a single AMD EPYC 9454 CPU. The GFN-FF_C_*_α_* optimisation for 4 structures did not converge, and for 15 structures it exceeded our operating memory limit of 196 GB. Therefore, only the remaining 461 protein structures were used for further analysis. All four datasets, along with the scripts used, are available in the supplementary files.

## 4 Comparison of PROPTIMUS RAPHAN_GFN-FF_ and GFN-FF_C*α*_ optimised structures

GFN-FF_C_*_α_* is a reference method to PROPTIMUS RAPHAN_GFN-FF_. Therefore, in this Section, SET_C_*_α_* with SET_RAPHAN_ are compared. SET_ORIG_ is also included in this comparison to evaluate differences between SET_C_*_α_* and SET_RAPHAN_ relative to the original structures (see Figure 2). The structures were compared in terms of the mean absolute deviation (MAD)s of atom positions, bond lengths, bond angles and dihedral angles. In addition, the MADs of canonical dihedral angles *ϕ*, *ψ*, *ω* and the side-chain dihedral angle *χ*_1_, acting as soft descriptors for local displacements in the protein backbone, were calculated [17]. The comparative analysis results are summarised in Table 1. Percentiles and associated histograms of differences are available in the supplementary files.

**Table 1.** Comparison of SET_C*α*_ with sets of structures SET_RAPHAN_ and SET_ORIG_. Mean absolute deviation of atom positions, bond lengths, bond angles, dihedral angles and four characteristic dihedral angles are included.

|  | SET<sub>RAPHAN</sub><br>vs.<br>SET<sub>Cα</sub> | SET<sub>ORIG</sub><br>vs.<br>SET<sub>Cα</sub> |
| --- | --- | --- |
| MAD atomic position [$\text{\AA}$] | 0.074 | 0.407 |
| MAD bond length [pm] | 0.075 | 4.851 |
| MAD bond angle [$^\circ$] | 0.136 | 2.071 |
| MAD dihedral angle [$^\circ$] | 1.611 | 11.124 |
| MAD $\phi$ [$^\circ$] | 1.814 | 10.275 |
| MAD $\psi$ [$^\circ$] | 1.686 | 9.227 |
| MAD $\omega$ [$^\circ$] | 1.156 | 5.770 |
| MAD $\chi_1$ [$^\circ$] | 1.307 | 7.981 |

The data in Table 1 show a high similarity between SET_RAPHAN_ and SET_C_*_α_*. The structures exhibit the highest degree of similarity in bond lengths, with a MAD of only 0.075 pm, which is at the level of accuracy of the PDB format. The MAD values for atomic positions (0.074 AÅ) and bond angles (0.136°) also show good agreement. The MADs for dihedral angles range around 1.5°. Relative to the MADs between SET_ORIG_ and SET_C_*_α_*, all these values are 5 *−* 65*×* lower, with the most significant improvement occurring in bond lengths and the smallest in dihedral angles.

### 4.1 Comparison of PROPTIMUS RAPHAN_GFN-FF_ optimised structures with their corresponding GFN-FF_C_*_α_* local minima

Based on the results in Table 1, it is not possible to determine to what extent the remaining differences between SET_RAPHAN_ and SET_C_*_α_* are due to the approximative nature of PROPTIMUS RAPHAN_GFN-FF_ (i.e., the neglect of long-range interactions) and how much they stem from PROPTIMUS RAPHAN_GFN-FF_ converging to a different local minima on the GFN-FF_C_*_α_* potential energy surface representing an alternative protein conformation. It is not at all unusual for various optimisation techniques to converge to more or less different minima in a complex space [32]. For this reason, three additional comparisons were performed (see Figure 2):

- The comparison of SET_RAPHAN_ and SET_RAPHAN+C_*_α_* evaluates the differences between structures optimised by PROPTIMUS RAPHAN_GFN-FF_ and their corresponding local minima on the potential energy surface reached by applying GFN-FF_C_*_α_*. Therefore, this comparison assesses the error caused by the approximative nature of PROPTIMUS RAPHAN_GFN-FF_relative to the nearest reachable local minima on the GFN-FF_C_*_α_* potential energy surface.
- The comparison of SET_C_*_α_* and SET_RAPHAN+C_*_α_* expresses the differences between local minima on the GFN-FF_C_*_α_* potential energy surface reached by applying GFN-FF_C_*_α_* on the original structures versus on structures optimised by the PROPTIMUS RAPHAN_GFN-FF_ method. This comparison detects whether PROPTIMUS RAPHAN_GFN-FF_ converges to a different local minimum on the GFN-FF_C_*_α_* potential energy surface than the original GFN-FF_C_*_α_* method does.
- The comparison of SET_ORIG_ and SET_RAPHAN+C_*_α_* allows an evaluation of the differences between SET_RAPHAN_ and SET_RAPHAN+C_*_α_* relative to the original structures.

The sets were compared using the same methodology as in Section 4. The comparative analysis results are summarised in Table 2. Percentiles and associated histograms of differences are available in the supplementary files.

**Table 2.** Comparison of SET_RAPHAN+C*α*_ with other sets of structures SET_C*α*_, SET_RAPHAN_ and SET_ORIG_. Mean absolute deviation of atom positions, bond lengths, bond angles, dihedral angles and four characteristic dihedral angles are included.

| | $\text{SET}_{\text{RAPHAN}}$<br>vs.<br>$\text{SET}_{\text{RAPHAN}+\text{C}\alpha}$ | $\text{SET}_{\text{C}\alpha}$<br>vs.<br>$\text{SET}_{\text{RAPHAN}+\text{C}\alpha}$ | $\text{SET}_{\text{ORIG}}$<br>vs.<br>$\text{SET}_{\text{RAPHAN}+\text{C}\alpha}$ |
| --- | --- | --- | --- |
| MAD atomic position [Å] | 0.033 | 0.056 | 0.412 |
| MAD bond length [pm] | 0.063 | 0.061 | 4.851 |
| MAD bond angle [°] | 0.080 | 0.105 | 2.071 |
| MAD dihedral angle [°] | 0.664 | 1.257 | 11.143 |
| MAD $\phi$ [°] | 0.881 | 1.339 | 10.302 |
| MAD $\psi$ [°] | 0.816 | 1.246 | 9.247 |
| MAD $\omega$ [°] | 0.596 | 0.839 | 5.778 |
| MAD $\chi_1$ [°] | 0.590 | 0.993 | 8.018 |

A comparison reveals that, for all tested metrics except bond lengths, the structures from SET_RAPHAN_ and SET_RAPHAN+C_*_α_* exhibit approximately half the difference compared to the structures from SET_RAPHAN_ and SET_C_*_α_* from Section 4. In addition, the comparison between SET_C_*_α_* and SET_RAPHAN+C_*_α_* proves the difference of these two sets of local minima on the GFN-FF_C_*_α_* potential energy surface. This confirms that the PROPTIMUS RAPHAN_GFN-FF_ and GFN-FF_C_*_α_* methods converge to different local minima on the GFN-FF_C_*_α_* potential energy surface. This fact is further demonstrated by the histogram of atomic position differences shown in Figure 3a). The histogram also highlights that the methods converge to alternative conformations, particularly for residues whose side chains form few or no hydrogen bonds (i.e., non-polar residues and residues on the structure’s surface). These regions are naturally flexible: the side chains switch between several energetically similar conformations.

**Fig. 3.**
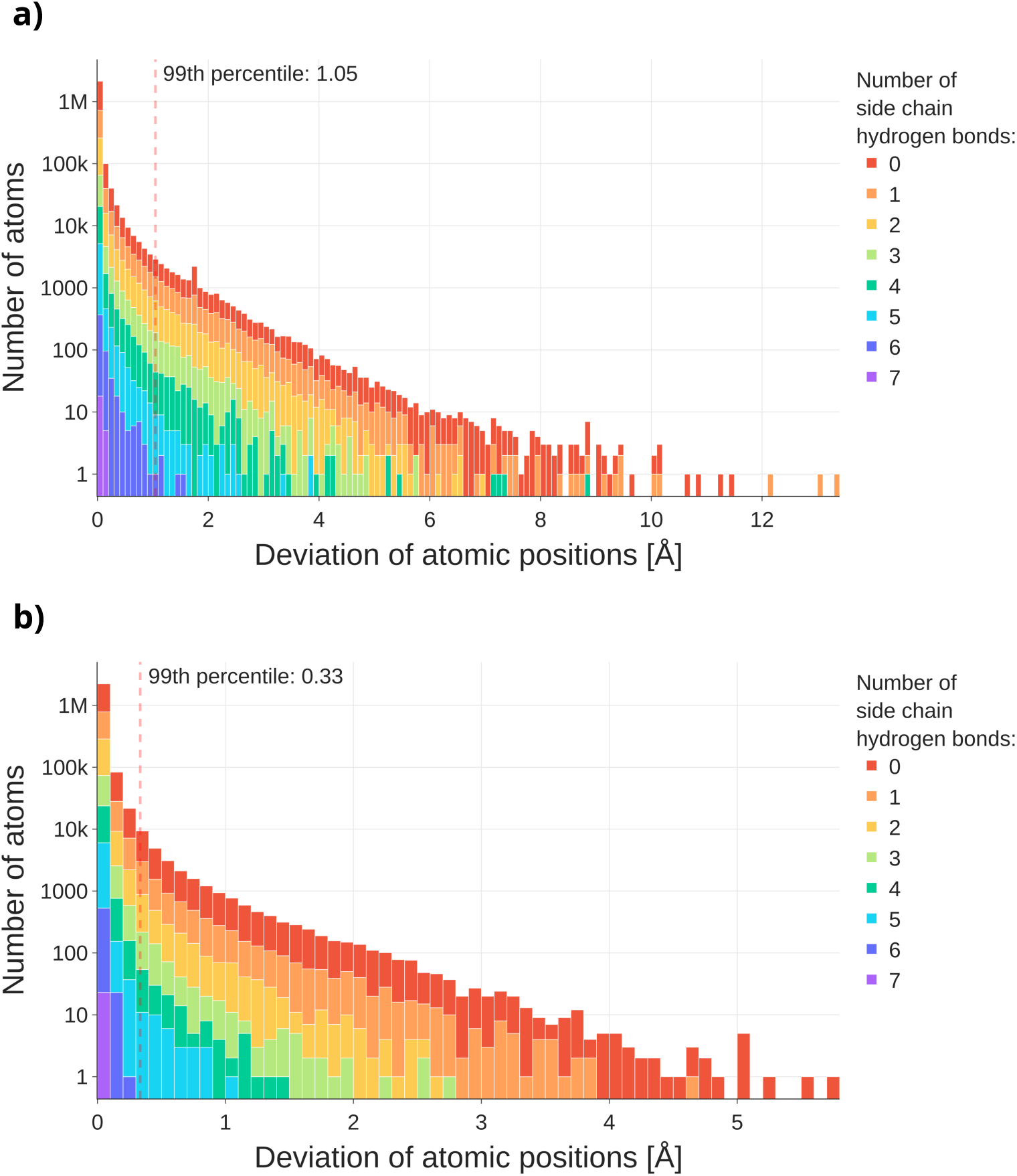
Histograms of atom position deviations between a) SET_C*α*_ and SET_RAPHAN+C*α*_ and b) SET_RAPHAN_ and SET_RAPHAN+C*α*_. Bars are coloured according to the number of hydrogen bonds of the side chain residue to which they belong. The image was created using the Plotly tool [33].

A comparison between the SET_RAPHAN_ and SET_RAPHAN+C_*_α_* sets in Table 2 shows that the atomic positions of protein structures optimised by PROPTIMUS RAPHAN_GFN-FF_ deviate, on average, by 0.033 AÅ from their positions in the corresponding reachable local minimum on the GFN-FF_C_*_α_* potential energy surface. The MAD of bond lengths is 0.063 pm. The MAD of bond angles is smaller than 0.1° and the one of dihedral angles is smaller than 1°. The most significant improvement relative to the original structures by the PROPTIMUS RAPHAN_GFN-FF_ optimisations is achieved in bond lengths and angles, where the MADs decrease by 77*×* and 26*×* respectively. The MADs for other similarity metrics decrease by 10 *−* 17*×*.

For visual comparison, the structure with the worst MAD of atomic positions between structures from SET_RAPHAN_ and SET_RAPHAN+C_*_α_* with less than a thousand atoms is shown in Figure 4. While most atoms are in good agreement, the atoms of residues CYS9 and GLY8 differ. To analyse such outliers between SET_RAPHAN_ and SET_RAPHAN+C_*_α_* in detail, a histogram of the atom position deviations distribution is visualised in Figure 3b). The histogram shows that on rare occasions, the deviations caused by the approximative nature of PROPTIMUS RAPHAN_GFN-FF_ can be very significant. Bars in the histogram are coloured according to the number of hydrogen bonds in the amino acid side chains. It can be seen, by examining the colours, that these rare significant deviations in atom positions are usually found within regions that are poor in hydrogen bonds, meaning nonpolar regions or regions on the surface of the structure. As previously mentioned, the side chains in these regions are naturally flexible and exhibit high conformational variability.

**Fig. 4.**
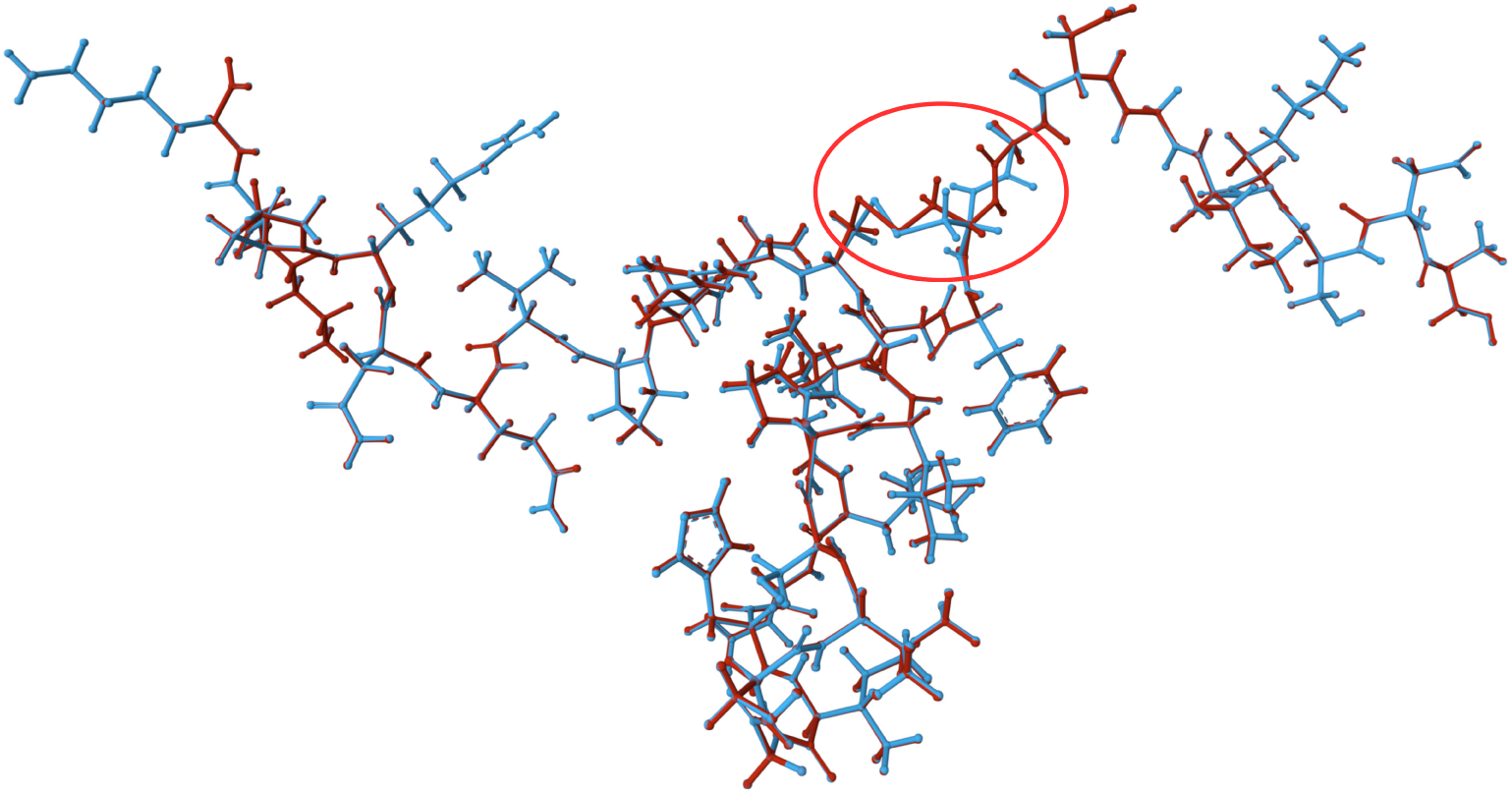
The structure with the worst MAD of atomic positions with less than a thousand atoms from the testing dataset with UniProtKB AC P83224 optimised by PROPTIMUS RAPHAN_GFN-FF_ (blue) and PROPTIMUS RAPHAN_GFN-FF_ + GFN-FF_C*α*_ (red). The MAD of atomic positions for this structure is 0.075 Å. While most atoms are in good agreement, the atoms of residues CYS9 and GLY8 differ (red circle). The Mol* visualisation software [34] was used to create the figure.

### 4.2 Comparison of optimisation times and memory usage

To evaluate the computational efficiency, the computational time of PROPTIMUS RAPHAN_GFN-FF_and GFN-FF_C_*_α_* optimisations of original structures was compared. This comparison is illustrated in Figure 5. In the case of GFN-FF_C_*_α_*, the computation time increases quadratically [17]. In contrast, the computation time of PROPTIMUS RAPHAN_GFN-FF_ increases linearly with respect to the number of atoms. The high variability of PROPTIMUS RAPHAN_GFN-FF_ calculation duration stems from the fact that the total runtime is also influenced by the number of residual substructures, their respective atom counts, and the number of iterations required for PROPTIMUS RAPHAN_GFN-FF_ convergence. The method achieves a minimum throughput of 3,600 atoms per hour and an average speed of 5,000 atoms per hour on the CPU used for testing.

In addition to computational speed, memory efficiency is a crucial factor for the accessibility of the optimisation process. As previously noted in Section 3.1, the GFN-FF_C_*_α_* optimisation of 15 structures exceeded the 196 GB memory limit. By contrast, PROPTIMUS RAPHAN_GFN-FF_ is significantly more memory-efficient. For the largest structure in our dataset (9,940 atoms), the optimisation required only 0.5 GB of RAM on a single CPU, and 3 GB when utilising 16 CPUs.

## 5 Conclusion

We developed the generic method PROPTIMUS RAPHAN as a fast alternative for the optimisation of proteins with constrained alpha carbons. Due to the use of the divide-and-conquer approach, the time complexity of PROPTIMUS RAPHAN is linear in relation to the size of the protein structure. We implemented our method as PROPTIMUS RAPHAN_GFN-FF_ employing the modern and almost QM-accurate force field GFN-FF and tested PROPTIMUS RAPHAN_GFN-FF_ on 461 structures from AlphaFold DB.

We found that the structures optimised by PROPTIMUS RAPHAN_GFN-FF_ are generally similar to those optimised by the GFN-FF_C_*_α_* method. However, the results also demonstrate that PROPTIMUS RAPHAN_GFN-FF_ converges to different local minima on the GFN-FF_C_*_α_* potential energy surface than the GFN-FF_C_*_α_* method itself, particularly in flexible regions with a low number of side-chain hydrogen bonds. On the other hand, the structures optimised by PROPTIMUS RAPHAN_GFN-FF_ are comparable to their corresponding reachable local minima on the GFN-FF_C_*_α_* potential energy surface, where the average atomic position deviation is 0.033 AÅ with outliers again occurring in flexible regions. Thus, while PROPTIMUS RAPHAN_GFN-FF_ may con-verge to different conformations than the original GFN-FF_C_*_α_* method, it approaches these alternative conformations with minimal error.

**Fig. 5.**
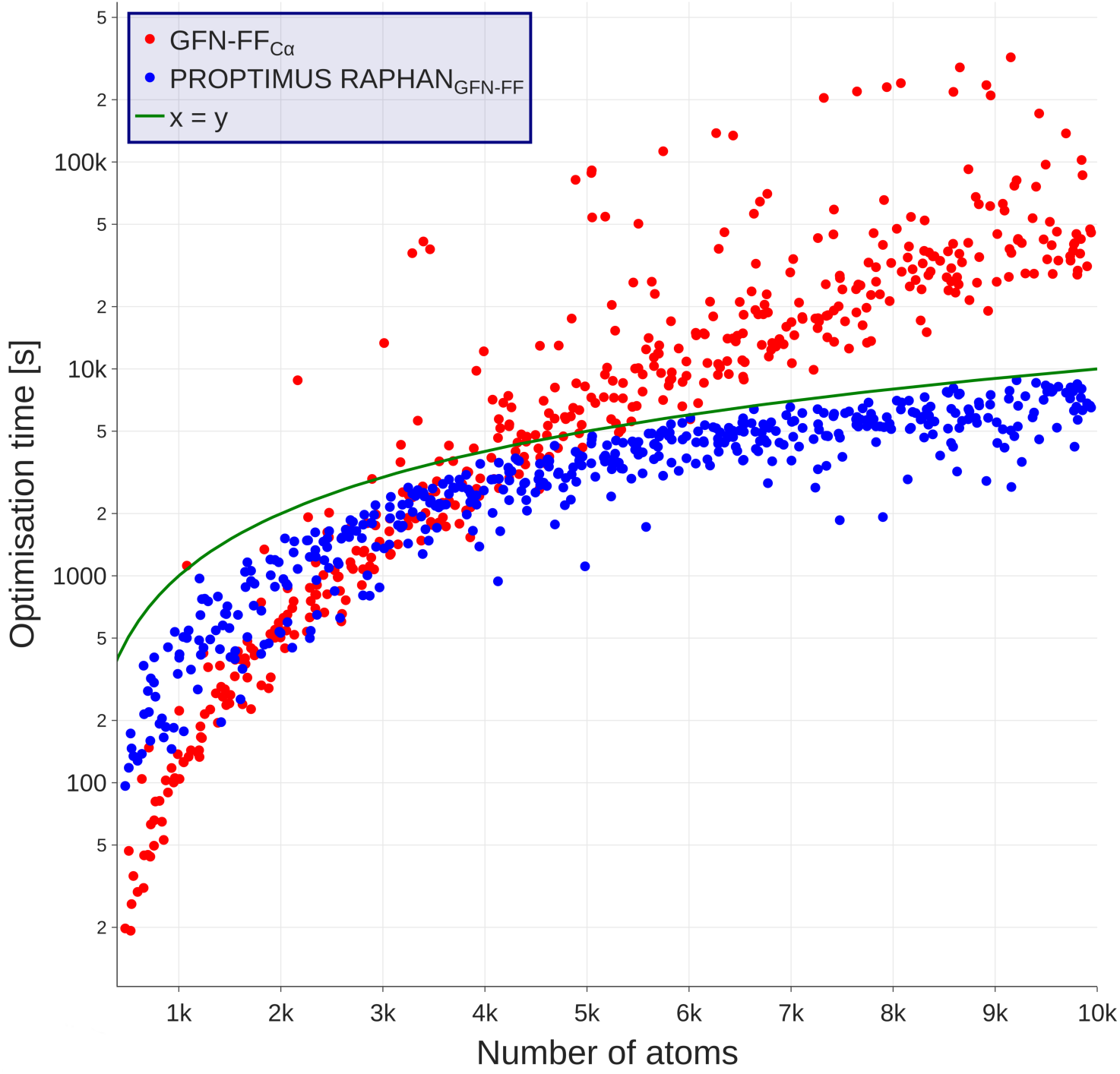
Comparison of computational times of PROPTIMUS RAPHAN_GFN-FF_ and GFN-FF_C*α*_ optimisations. The image was created using the Plotly tool [33].

While the computation time for optimisation with constrained alpha carbons increases quadratically, the average speed of PROPTIMUS RAPHAN_GFN-FF_ is 5 000 atoms per hour on the CPU used for testing purposes. Another advantage of PROPTIMUS RAPHAN_GFN-FF_ is its low RAM usage, which further enhances its accessibility and practical utility. PROPTIMUS RAPHAN_GFN-FF_ is freely available, parallelisable, easy to install, and simple to use. Even the largest structures in AlphaFold DB could be highly optimised by PROPTIMUS RAPHAN_GFN-FF_ on a standard desktop computer in several hours.

## Acronyms

*GFN-FF_Cα_* GFN-FF optimisation with constrained alpha carbons.

*MAD* mean absolute deviation.

*PROPTIMUS RAPHAN* Per-residue optimisation of protein structures: Rapid alternative to optimisation with constrained alpha carbons.

*PROPTIMUS RAPHAN_GFN-FF_* PROPTIMUS RAPHAN method employing a force field, GFN-FF.

*PROPTIMUS RAPHAN_GFN-FF_ + GFN-FF_Cα_* PROPTIMUS RAPHAN_GFN-FF_ method, followed by optimisation by the GFN-FF_C_*_α_* method.

## 6 Declarations

### 6.1 Availability of Data and Materials

The implementation of PROPTIMUS RAPHAN, including a description of its installation and use, is available at GitHub under the MIT licence at https://github.com/sb-ncbr/proptimus_raphan. The supplementary files supporting the conclusions of this article are available in the OneData repository at https://doi.org/10.58074/kap4-ka76 as well as archived implementation of PROPTIMUS RAPHAN.

### 6.4 Authors’ Contributions

Conceptualisation: OS. Data curation: OS, TS, GB. Formal analysis: OS, TS, GB. Funding acquisition: OS, RS. Investigation: OS, GB. Methodology: OS. Project administration: OS, GB, RS. Resources: OS, TS, RS. Software: OS, TS. Supervision: OS, RS. Validation: OS, TS, GB. Visualisation: OS, GB. Writing - original draft: OS, GB. Writing - review and editing: All authors.

## Acknowledgements

Biological Data Management and Analysis Core Facility of CEITEC Masaryk University, funded by ELIXIR CZ research infrastructure (MEYS Grant No: LM2023055), is gratefully acknowledged for supporting the research presented in this paper. Computational resources were provided by the e-INFRA CZ (ID: 90254). The authors thank Lukáš Bohuš for valuable assistance with the manuscript review.

